# Auranofin induces disulfide bond-mimicking S-Au-S bonds in protein thiol pairs

**DOI:** 10.1101/2024.01.15.575694

**Authors:** Laísa Quadros Barsé, Petra Düchting, Natalie Lupilov, Julia E. Bandow, Ute Krämer, Lars I. Leichert

## Abstract

Auranofin is an inhibitor of human thioredoxin reductase, clinically used in the treatment of rheumatoid arthritis. More recently, it has been shown to possess strong antibacterial activity. Despite the structural dissimilarity and the independent evolutionary origins of human thioredoxin reductase and its bacterial counterpart (TrxB), inhibition of bacterial thioredoxin reductase is often suggested to be a major factor in auranofin’s antibacterial mode of action. To test this hypothesis, we attempted to determine the mechanism of inhibition of auranofin for bacterial TrxB in the presence of thioredoxin, TrxB’s natural substrate. However, the data obtained in these experiments was not consistent with a specific and exclusive interaction between TrxB and auranofin. Instead, it suggested that auranofin directly interacts with the cysteine thiols in thioredoxin, TrxB’s substrate. Using the fluorescent redox protein roGFP2, we showed that auranofin does indeed directly interact with cysteine pairs in proteins, forming a thiol modification that is similar to, but clearly distinct from a disulfide bond. The Au:S stoichiometries of auranofin-treated roGFP2 and thioredoxin strongly suggest the presence of an S-Au-S bridge between two cysteines in those proteins. These S-Au-S bonds form independent of thioredoxin reductase at a rate that indicates their pertinence in auranofin’s antibacterial mode of action.

## Introduction

Auranofin is a gold(I)-containing drug developed for the treatment of rheumatoid arthritis (Finkelstein et al., 1976). Its mode of action is the inhibition of thioredoxin reductase (TrxR), the NADPH-dependent part of the thioredoxin thiol-disulfide oxidoreductase system (Hwang-Bo et al., 2017; Rigobello et al., 2005; Zhang et al., 2019). Thioredoxin reductase inhibition is mediated through direct binding of gold(I) to a crucial selenocysteine in the enzyme’s C-terminus (Pickering et al., 2020). From observations made in the related enzyme thioredoxin glutaredoxin reductase (TGR) from *Schistosoma mansoni*, it has been suggested that an Se-Au-S bond between a selenocysteine and cysteine is initially formed in the C-terminus and that the gold can then be transferred to cysteine pairs within the enzyme, including the active site cysteine pair close to the FAD-cofactor binding site (Saccoccia et al., 2012, 2014).

Recent studies have shown that auranofin has an additional strong antibiotic activity against pathogenic bacteria, including *E. faecalis, H. pylori, M. tuberculosis,* and multi-drug resistant *S. aureus* (Cassetta et al., 2014; Epstein et al., 2019; Harbut et al., 2015; Owings et al., 2016; She et al., 2019; Thangamani et al., 2016; Tharmalingam et al., 2019). It is tempting to hypothesize that auranofin’s antibacterial mode of action is, similar to its antirheumatic action, the inhibition of bacterial thioredoxin reductase (TrxB). Indeed, several studies show that bacterial thioredoxin reductase is inhibited effectively by auranofin at nanomolar concentrations (Epstein et al., 2019; Harbut et al., 2015; Owings et al., 2016; Tharmalingam et al., 2023), and auranofin treatment induces the oxidative stress response in both Gram-positive and Gram-negative species (Barse et al., 2023; Senges et al., 2020), an observation seemingly indicative of the inhibition of the main cellular thiol-disulfide oxidoreductase system.

However, mammalian TrxR and bacterial TrxB are structurally highly distinct and are thought to have evolved independently (Lu & Holmgren, 2014b, 2014a), with the bacterial enzyme lacking the selenocysteine-containing domain that is directly involved in the initial gold binding in the mammalian enzyme (Biterova et al., 2005; Cheng et al., 2009; Pickering et al., 2020; Sandalova et al., 2001; Waksman et al., 1994; Williams, 1995). Thus, if inhibition of bacterial thioredoxin reductase is a major factor in auranofin’s antibacterial activity, its mechanism of inhibition must be distinct from the mechanism of inhibition of mammalian thioredoxin reductase.

Here we show that, in the presence of its natural substrate thioredoxin (TrxA), bacterial thioredoxin reductase TrxB is not effectively inhibited by auranofin. Using the engineered redox probe protein roGFP2, we demonstrate that auranofin directly interacts with thiol pairs in proteins in a way that mimics a disulfide bond. Determination of the stoichiometry of gold binding with ICP-OES in auranofin-treated roGFP2 and TrxA suggests the presence of S-Au-S bridges in these proteins.

## Materials & Methods

### Construction of plasmids encoding components of the bacterial Trx system

Bacterial strains, plasmids, and oligonucleotides used in this study are listed in Table 1. Plasmid construction followed standard procedures. Newly constructed plasmids were verified by DNA sequencing.

**Table 1:**
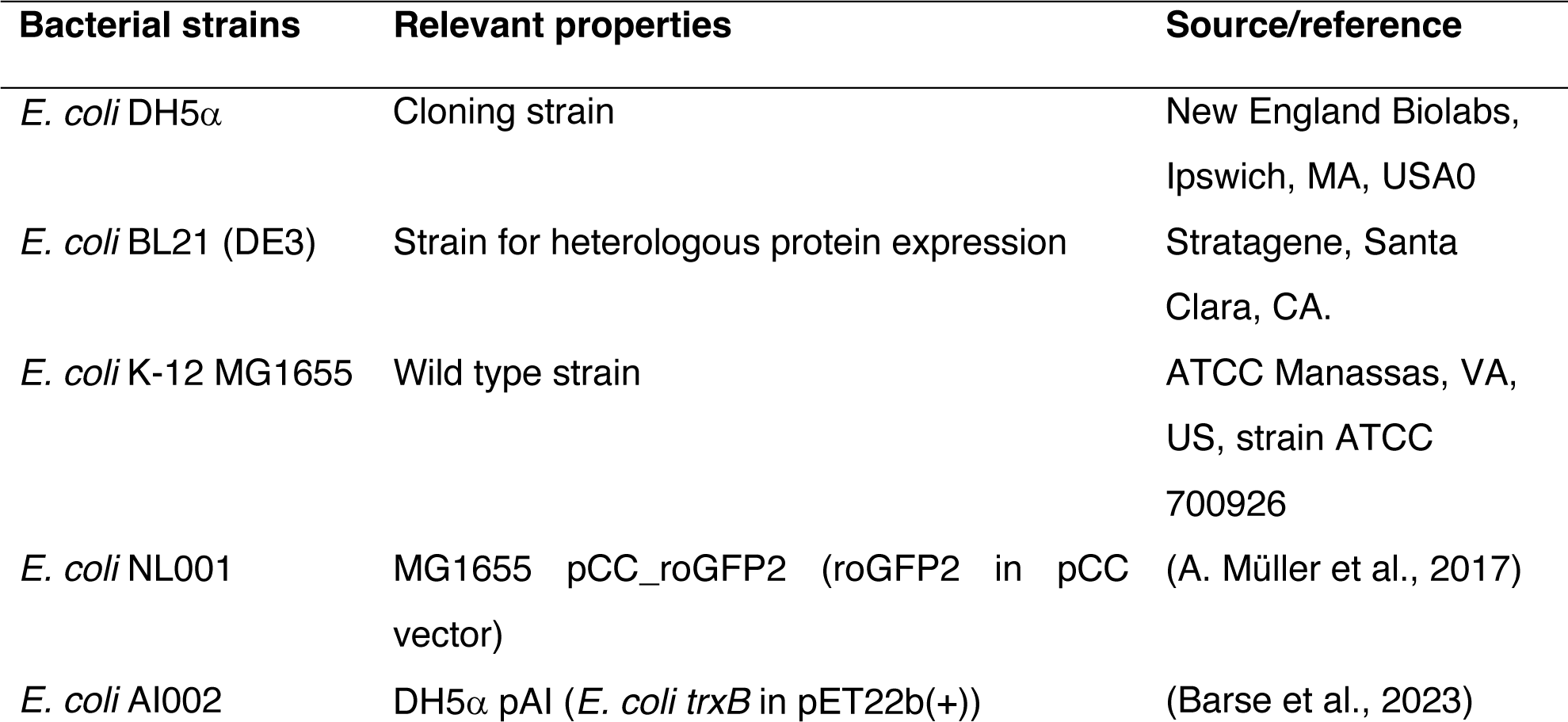
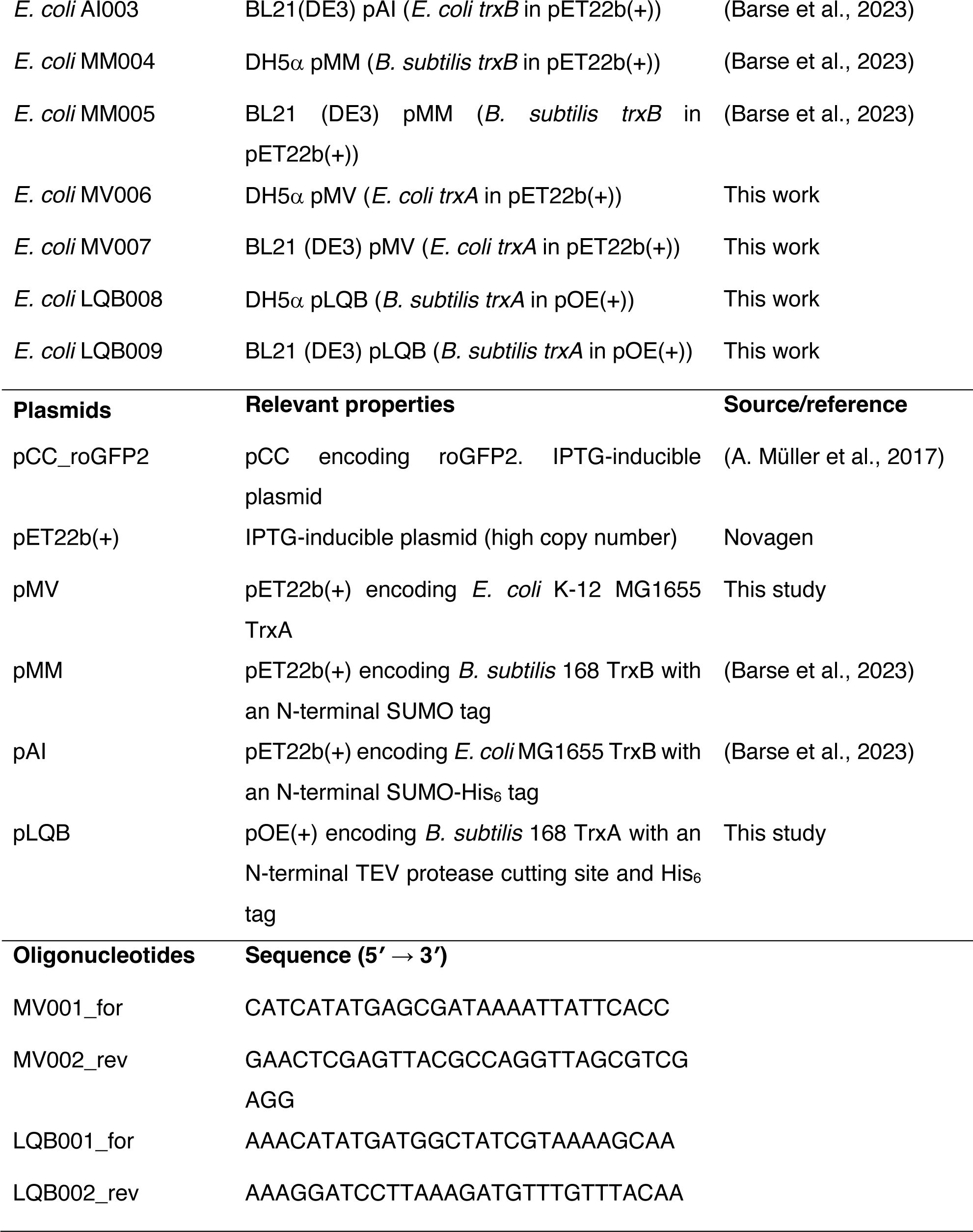
Bacterial strains, plasmids, and oligonucleotides used in this study.

The *trxB* genes encoding *E. coli* MG1655 TrxB and *B. subtilis* 168 TrxB were cloned into pET22b(+) vectors, fused to a SUMO-tag, as previously described (Barse et al., 2023) yielding plasmids pAI and pMM.

The *trxA* gene from *E. coli* MG1655 encoding thioredoxin was amplified from *E. coli* MG1655 genomic DNA via PCR with the primers MV001_for and MV002_rev. The PCR product was digested with NdeI and XhoI and cloned into the pET22b(+) vector, yielding the plasmid pMV.

The *trxA* gene from *B. subtilis* encoding thioredoxin was amplified from *B. subtilis* 168 genomic DNA via PCR with the primers LQB001_for and LQB002_rev. The PCR product was digested with NdeI and BamHI and cloned into the pOE vector (Nilewski et al., 2021), yielding the plasmid pLQB, containing *trxA* fused to an N-terminal His_6_-tag and a TEV protease cutting site.

### Heterologous protein expression

For heterologous protein expression, a single colony of *E. coli* MV007, or LQB009 carrying pMV, or pLQB was used to inoculate 50 mL LB containing 200 mg/L ampicillin, and the cells were grown overnight at 37 °C and 120 rpm. The overnight culture was then used to inoculate 5 L of LB medium supplemented with ampicillin. Cells were incubated at 37 °C and 120 rpm until an OD_600_ of 0.5-0.6 was reached. Subsequently, protein expression was induced by adding 1 mM isopropyl 1-thio-ß-D-galactopyranoside (IPTG) to the cell culture. After overnight incubation at 20 °C and 120 rpm, cells were harvested by centrifugation at 7800 x *g* and 4 °C for 45 min, and pellets were stored at −80 °C.

Heterologous roGFP2 expression and purification, as well as *E. coli* and *B. subtilis* TrxB expression and purification, followed established protocols (Barse et al., 2023; A. Müller et al., 2017).

### Purification of thioredoxins

The cell pellet obtained after TrxA overexpression in *E. coli* MV007 (*ec*TrxA) was washed once with 80 mL of lysis buffer 1 (20 mM Tris-HCl, pH 8.0) containing 2 mL of EDTA-free protease inhibitor mixture (Roche Applied Science, Penzberg, Germany). Cells were disrupted by passing the cell suspension through a Constant Cell disruption system (TS benchtop; Constant Systems, Daventry, UK) three times at 1.9 kbar and 4 °C, followed by the addition of the serine protease inhibitor phenylmethylsulphonyl fluoride (PMSF) to a final concentration of 1 mM. The cell lysate was centrifuged at 6700 x *g* and 4 °C for 1 h. The supernatant was vacuum-filtered through a 0.45 μm filter. Purification and fractionation steps were performed with an “ÄKTApurifier” FPLC system (GE Healthcare). The filtrate was loaded onto a HiTrap DEAE FF column (5 mL, Cytiva) pre-equilibrated with lysis buffer 1.

Subsequently, the column was washed with 150 mL lysis buffer 1, and then eluted with a gradient (2-100 %) of elution buffer 1 (40 column volumes, 20 mM Tris-HCl, 1M NaCl, pH 8.0). 3 mL fractions were collected and analyzed by SDS-PAGE. The combined elution fractions containing *ec*TrxA were concentrated to 5 ml using a Vivaspin 20 PES, MWCO 5000 concentrator system (Sartorius Stedim Biotech). The protein solution was then loaded onto a size exclusion Superdex 75 26/600 column (Cytiva), pre-equilibrated with 1 column volume of 40 mM sodium phosphate, 200 M NaCl, pH 7.5. Fractions containing purified *ec*TrxA were pooled and concentrated using the Vivaspin 20 PES, MWCO 5000 concentrator system. Purified *ec*TrxA was stored in sodium phosphate buffer supplemented with 10 % (v/v) glycerol at −80 °C.

*E. coli* LQB009 cell pellets obtained after overexpression of *B. subtilis* 168 TrxA (*bs*TrxA) were resuspended in 30 mL lysis buffer 2 (50 mM sodium phosphate, 300 mM NaCl, pH 8.0) containing 2 mL of EDTA-free protease inhibitor mixture. Cell disruption and preparation of the crude protein extract followed the steps outlined for *ec*TrxB. The resulting filtrate was loaded onto 2 × 5 ml nickel–nitrilotriacetic acid (Ni–NTA) columns “HisTrapTM HP” (GE Healthcare) equilibrated with lysis buffer 2. Elution was performed with 5 column volumes of elution buffer 2 (lysis buffer 2 containing 500 mM imidazole) in a gradient (2-100%). 3 mL fractions were collected and analyzed by SDS-PAGE. Fractions with the highest target protein yield were pooled, and His_6_-tagged TEV protease was added (30-fold excess of the target protein to TEV protease). The protein solution was dialyzed against 5 L TEV buffer (50 mM Tris-HCl pH 8, 1 mM EDTA) in dialysis tubes with an 8 kDa cutoff, overnight at 4 °C. Then, 2 mM MgCl_2_ was added to the pooled protein, and the protein solution was loaded again onto the Ni–NTA columns to remove the His_6_-tagged TEV protease, as well as the cleaved His_6_ tag. Flowthrough containing the protein was dialyzed against 100 mM sodium phosphate buffer (pH 7.0) containing 1 mM EDTA. The protein was concentrated using the Vivaspin 20 PES, MWCO 5000 concentrator system. Purified *bs*TrxA was stored in sodium phosphate buffer supplemented with 10 % (v/v) glycerol at −80 °C.

### Reconstitution of the bacterial thioredoxin system and inhibition assays

TrxB activity in the absence or presence of auranofin was determined by performing a 5,5’-dithiobis(2-nitrobenzoic) acid (DTNB) reduction assay. In this assay, the NADPH-dependent TrxB reduces oxidized TrxA, which then reacts with DTNB to yield the spectrophotometrically detectable 5-thio 2-nitrobenzoic acid (TNB). DTNB assays were performed in transparent, flat-bottom 96-well plates (Sarstedt, Nümbrecht, Germany) as described by Lu *et al*. (Lu et al. 2013). Briefly, 50 nM *ec*TrxB or *bs*TrxB were pre-incubated for 10 minutes with various amounts of TrxA and auranofin (10 mg/mL stock in DMSO) in 100 µL TE buffer (50 mM Tris-HCl, 2 mM EDTA, pH 7.5) containing 200 µM NADPH (Sigma-Aldrich, St. Louis, USA). Protein concentration was determined through absorption at 280 nm using the molar extinction coefficients 18910 M^-1^cm^-1^ (*ec*TrxB), 24870 M^-1^cm^-1^ (*bs*TrxB), 13980 M^-1^cm^-1^ (ecTrxA), and 12490 M^-1^cm^-1^ (bsTrxA). Extinction coefficients were calculated by ProtParam (Gasteiger et al., 2005). The enzymatic reaction was initiated by adding 100 µL TE buffer containing 200 µM NADPH and 5 mM DTNB (Sigma-Aldrich, St. Louis, USA). The amount of liberated TNB was then quantified spectrophotometrically by measuring the absorbance at 412 nm and using an extinction coefficient of 13600 M^−1^cm^−1^.

Initial reaction rates (V_0_; DTNB [µM]/min) of *ec*TrxB and *bs*TrxB without auranofin were calculated from data points collected during the first 2 minutes after DTNB addition using linear regression and the trendline function in Excel. The initial velocities were then plotted against the TrxA concentration as a best-fit line using non-linear regression of the Michaelis-Menten equation (1) and the GraphPad Prism software. Assays were performed in triplicates.

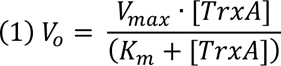

For the underlying kinetics data refer to supplementary material S1.

### TrxA inhibition in an insulin reduction assay

The insulin reduction assay was performed as described by Nilewski *et al*. (Nilewski et al., 2021). Briefly, FITC-labeled insulin reduction was monitored fluorometrically with a fluorescence spectrophotometer FP-8500 (Jasco) equipped with a Peltier-thermo cell holder EHC-813. Emission was followed over time at a wavelength of 520 nm at an excitation wavelength of 490 nm. Emission and excitation bandwidth was 2.5 nm, sensitivity was set to low, and one measurement was performed per second. Samples were prepared in a Quartz Suprasil cuvette for fluorescence measurements (Hellma). This experiment was done as a one-reaction cycle experiment without an additional reductant driving catalysis, so TrxA was pre-reduced using 1 mM dithiothreitol (DTT) (Sigma-Aldrich, St. Louis, USA), and DTT was subsequently removed using a “Micro Bio-Spin® Columns with Bio-Gel® P-30” (Bio-Rad).

The protein concentration was determined by absorption at 280, as described above. The assay buffer contained 20 mM potassium phosphate (pH 7.0), 2 mM EDTA, and 10 μg/ml FITC-labeled insulin (Sigma-Aldrich), and the fluorescence was measured at 25 °C while stirring (800 rpm). After stabilization of the fluorescent signal, *ec*TrxA or *bs*TrxA were added to a final concentration of 1 μM. In samples in which auranofin was included, the stated amount of auranofin was added right after the addition of thioredoxin from a 10 mg/mL stock in DMSO.

The maximum slope of fluorescence increase after the addition of thioredoxin over 250 s was determined with the Microsoft Excel software’s SLOPE and MAX functions and set as the respective velocity and expressed as Δa.u./min. Significance was calculated using Excel’s TTEST function assuming one-tailed distribution and two sample, equal variance. See supplementary material S2.

### Measurement of the oxidation degree of the roGFP2 probe *in vitro* in the presence of auranofin

Purified roGFP2 protein concentration was determined through absorption at 280 nm using its calculated molar extinction coefficient (21890 M^-1^cm^-1^), and it was diluted to 0.2 µM in PBS pH 7.4 (Gibco) + 5 mM EDTA buffer. Fluorescence measurements were performed in a JASCO FP-8500 fluorescence spectrometer equipped with a Peltier thermo-holder ‘EHC-813’ at 25 °C under continuous stirring (800 rpm). Measurement parameters were set to 510 nm (Em), 350–500 nm (Ex), 5 nm slit width (Ex/Em), and medium sensitivity. Samples were prepared in a Quartz Suprasil cuvette for fluorescence measurements (Hellma). Spectral scans were recorded in 1 min steps. After the first measurements, auranofin (10 mg/mL stock in DMSO) was added to a final concentration of 2 µM. After 15 min, DTT was added at a final concentration of 20 µM to fully reduce the probe.

The ratio of the fluorescence excitation intensities (405/488 nm) was used to calculate the probe’s oxidation state. All values were normalized to OxD using fully oxidized (AT-2-treated) and fully reduced (DTT-treated) roGFP2 and the following equation (2):

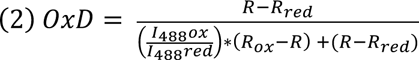

*R_ox_* is the 405/488 ratio of oxidized (AT-2-treated) and *R_red_* of reduced (DTT-treated) roGFP2, respectively. *I*_488_*ox* and *I*_488_*red* are the fluorescence intensities of roGFP2 at 488 nm under oxidizing or reducing conditions. *R* is the measured 405/488 nm ratio of roGFP2.

For calculation and normalization of the probe’s oxidation degree at each time point, the means of values recorded over time for *R_red_*, *R_ox_*, *I*_488_*ox*, and *I*_488_*red* were used. Data was processed using Microsoft Excel software and GraphPad Prism. See supplementary material S3 for the underlying data.

### Quantitation of Au using inductively coupled plasma-optical emission spectrometry (ICP-OES)

The elemental concentration of Au was quantified by Inductively Coupled Plasma Optical Emission Spectrometry (ICP-OES) using an iCAP 6500 Duo View ICP Spectrometer (Thermo Fisher Scientific, Waltham, USA) against a dilution series of auranofin used as calibration standard (Sigma-Aldrich, St. Louis, USA). roGFP2 and *ec*TrxA had their concentration determined by A_280_ absorption as outlined above, and then a 10-fold molar excess of DTT was added to fully reduce the protein. Proteins were incubated for 10 minutes at RT. DTT was removed using “Micro Bio-Spin® Columns with Bio-Gel® P-30” (Bio-Rad).

After DTT removal, a 10-fold molar excess of auranofin or 1000-fold molar excess of iodoacetamide was added to reduced roGFP2 or TrxA in PBS pH 7.4 (Gibco) + 5 mM EDTA. The samples were incubated on a ThermoMixer (Eppendorf, Hamburg, Germany) at RT and 800 rpm. After incubation for 1 hour, iodoacetamide or auranofin were added to the samples where appropriate, and excess amounts of auranofin were removed using “Micro Bio-Spin® Columns with Bio-Gel® P-30” (Bio-Rad). Additionally, buffer without protein but with auranofin was subjected to auranofin-removal by “Micro Bio-Spin® Columns with Bio-Gel® P-30” to control for carryover of auranofin in this procedure. The eluted protein concentration was again measured by A_280_ absorption. Appropriate sample volumes ranging from 50 µl to 75 µl to adjust the final protein concentration in the 10 mL analyte solution to 350 nM for roGFP2 and 200 nM for TrxA, were digested by addition of 1 mL 65% w/w nitric acid (Bernd Kraft) and heating to 80°C for 16 h, followed by the addition of ultrapure water to a final volume of 10 mL. The calibration curve was prepared using the following final concentrations of auranofin: 0, 62.5, 125, 250, 500, and 1000 nM. Calibrations standards were processed as described for the protein samples. Emission wavelengths were recorded at 242.795 nm. Additional data for 267.595 nm was collected to control for potential interference and this data was in agreement with the data reported (see also supplementary material S4). Each sample was quantified three times, and the average value was used. Three or two independently prepared replicates were measured for each sample, the acid blank control was measured only once.

### Removal of Au from roGFP2 by TrxA

Fluorescence spectrometry was used to measure the removal of Au from auranofin-treated roGFP2 by TrxA. Briefly, PBS (pH 7.4) (Gibco) + 5 mM EDTA was used as a buffer, then 100 µM roGFP2 was treated with a 10-fold molar excess of auranofin for 20 minutes. Excess auranofin was removed using “Micro Bio-Spin® Columns with Bio-Gel® P-30” (Bio-Rad) pre-equilibrated with PBS (pH 7.4) (Gibco) + 5 mM EDTA. The protein concentration was determined by A_280_ absorption and adjusted to 200 nM roGFP2. 1 mL roGFP2 solution was then measured as described above. After the fluorescent signal was stable, TrxA was added at various concentrations ranging from a 5 to 40-fold molar excess (1 µM to 8 µM), and samples of auranofin-treated roGFP2 were measured for 1 h. A control sample of DTT-treated roGFP2 was also measured for 30 minutes. Experiments were done in duplicates. Calculation of the probe’s OxD were performed as described above. See supplementary material S5 for the underlying data.

## RESULTS

### In a reconstituted bacterial thioredoxin system auranofin is not a specific inhibitor of thioredoxin reductase

Thioredoxin reductase (TrxR) is the target of auranofin in mammalian cells (Gromer et al., 1998). Similarly, bacterial thioredoxin reductase (TrxB) is presumed to be a target, and TrxB’s inhibition is thought to be the major cause of its antibacterial activity (Harbut et al., 2015). TrxBs from Gram-positive species are inhibited by auranofin at micro-molar levels of the inhibitor *in vitro* (Epstein et al., 2019; Harbut et al., 2015; Tharmalingam et al., 2023).

We, as well, have recently shown that TrxBs of both Gram-negative and Gram-positive bacteria are already fully inhibited by stochiometric amounts of auranofin (Barse et al., 2023). However, in this assay, we used DTNB, a relatively poor substrate of bacterial TrxB (S. Müller et al., 1996; Susanti et al., 2017). Thus, to learn more about the nature of the inhibition, we revisited our TrxB activity assay, this time using its native substrate bacterial thioredoxin (TrxA). Our enzymatic reactions contained TrxB at 50 nM, NADPH at 400 µM, well above the reported K_m_ of TrxB for NADPH at 1.2 - 2.69 µM to ensure pseudo-first order reaction conditions (Lennon & Williams, 1997; Prongay et al., 1989; Williams, 1976), DTNB at 2 mM and TrxA at concentrations ranging from 0 to 6 µM. Activity, as measured by TNB release from DTNB, was dependent on the concentration of TrxA, and we observed a K_m_ of TrxB for TrxA of 1.35 ± 0.23 µM in the reconstituted *E. coli* thioredoxin system and a K_m_ of 1.72 ± 0.16 µM of TrxB for TrxA in the reconstituted *B. subtilis* system (Fig. 1).

**Figure 1.**
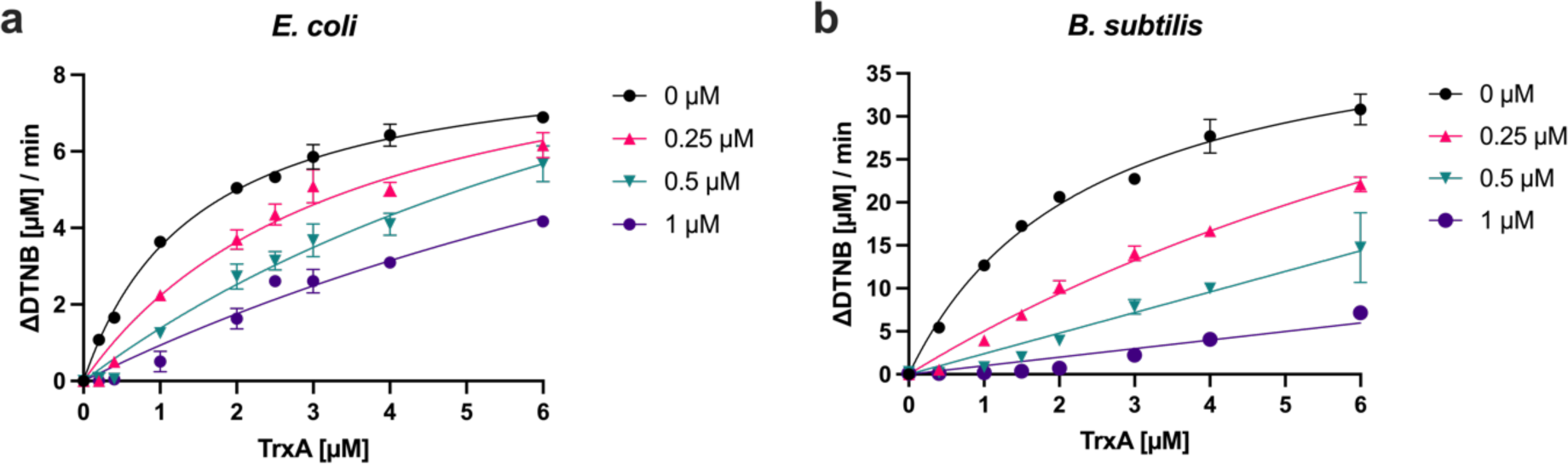
Auranofin inhibition of bacterial TrxB is attenuated by thioredoxin. Purified *ec*TrxB or *bs*TrxB (50 nM) were preincubated with 200 µM NADPH and increasing concentrations of *ec*TrxA or *bs*TrxA in the absence (0 µM) or presence of various concentrations of auranofin (0.25 - 1 µM). Reactions were initiated by adding 2 mM DTNB and additional 200 µM NADPH. Reaction progress was monitored by measuring the absorbance of the liberated TNB chromophore at 412 nm. Michaelis-Menten plots [initial velocity of reaction (DTNB [µM]/min) vs. substrate concentration (TrxA [µM])] of **(a)** *ec*TrxB and **(b)** *bs*TrxB activity are shown. Attempted fittings of the data for kinetics measured in the presence of auranofin are included to illustrate non-Michaelis-Menten behavior in the presence of inhibitor. The average and standard deviation of three independent experiments is shown. For the underlying data refer to supplementary material S1.

Next, we wanted to determine the inhibitor constant for auranofin for *ec*TrxB (Fig. 1, a) and *bs*TrxB (Fig. 1, b) by adding varying concentrations of auranofin to our enzymatic assay. However, this inhibition assay did not follow any pattern that would be expected if thioredoxin reductase was the only target of auranofin in our assay. In our previous assay, in the absence of thioredoxin, we saw full inhibition of TrxB already at equimolar concentrations of the inhibitor (Barse et al., 2023). However, now, in the presence of thioredoxin, we had to use concentrations substantially above the equimolar (50 nM TrxB) concentration to observe meaningful enzyme inhibition, especially at higher concentrations of thioredoxin. Additionally, any time we reached auranofin concentrations above the concentration of the substrate thioredoxin present in a given reaction, we achieved almost complete inhibition of TrxB activity. Inspection of our inhibition curves in Fig. 1, panels a and b, reveals that, at a concentration of 1 µM auranofin, discernible activity is only observed at TrxA concentrations at or above 1 µM, and similarly at concentrations of 0.25 µM and 0.5 µM auranofin and TrxA concentrations at or above 0.25 µM and 0.5 µM, respectively.

Based on these observations, we suspected that auranofin is binding to TrxA directly, as has been suggested by others (Saccoccia et al., 2014). Due to TrxA’s relatively high concentration as substrate in comparison to TrxB, it then attenuated the inhibition of the catalyst in a stoichiometric manner. Due to this unexpected behavior of auranofin, we could not deduce an inhibitor constant or a mechanism of inhibition from this data and had to dismiss our initial hypothesis that auranofin is a specific inhibitor of bacterial TrxB.

### Auranofin inhibits thioredoxin directly

To test if auranofin interacts with TrxA directly, we used TrxA from *E. coli* and *B. subtilis* in a thiol-disulfide oxidoreductase assay. Thioredoxin is known to facilitate the reduction of insulin (Holmgren, 1979). We used a variant of this classical insulin reduction assay (Nilewski et al., 2021) to check this inherent activity of TrxA in the presence of auranofin. In our assay, *ec*TrxA or *bs*TrxA were added as the reducing agent, and the velocity of insulin reduction was monitored spectrofluorometrically (Fig. 2). Adding stoichiometric amounts of auranofin led to a significant attenuation of insulin reduction. *ec*TrxA treated with auranofin at equimolar concentrations was 51 % slower than untreated *ec*TrxA, while autranofin-treated *bs*TrxA was 74.6 % slower (Fig. 2). These results corroborate our previous observation, and both results combined suggest that direct binding of auranofin or gold(I) to thioredoxin is the cause of the attenuation of the inhibition of bacterial thioredoxin reductase TrxB.

**Figure 2.**
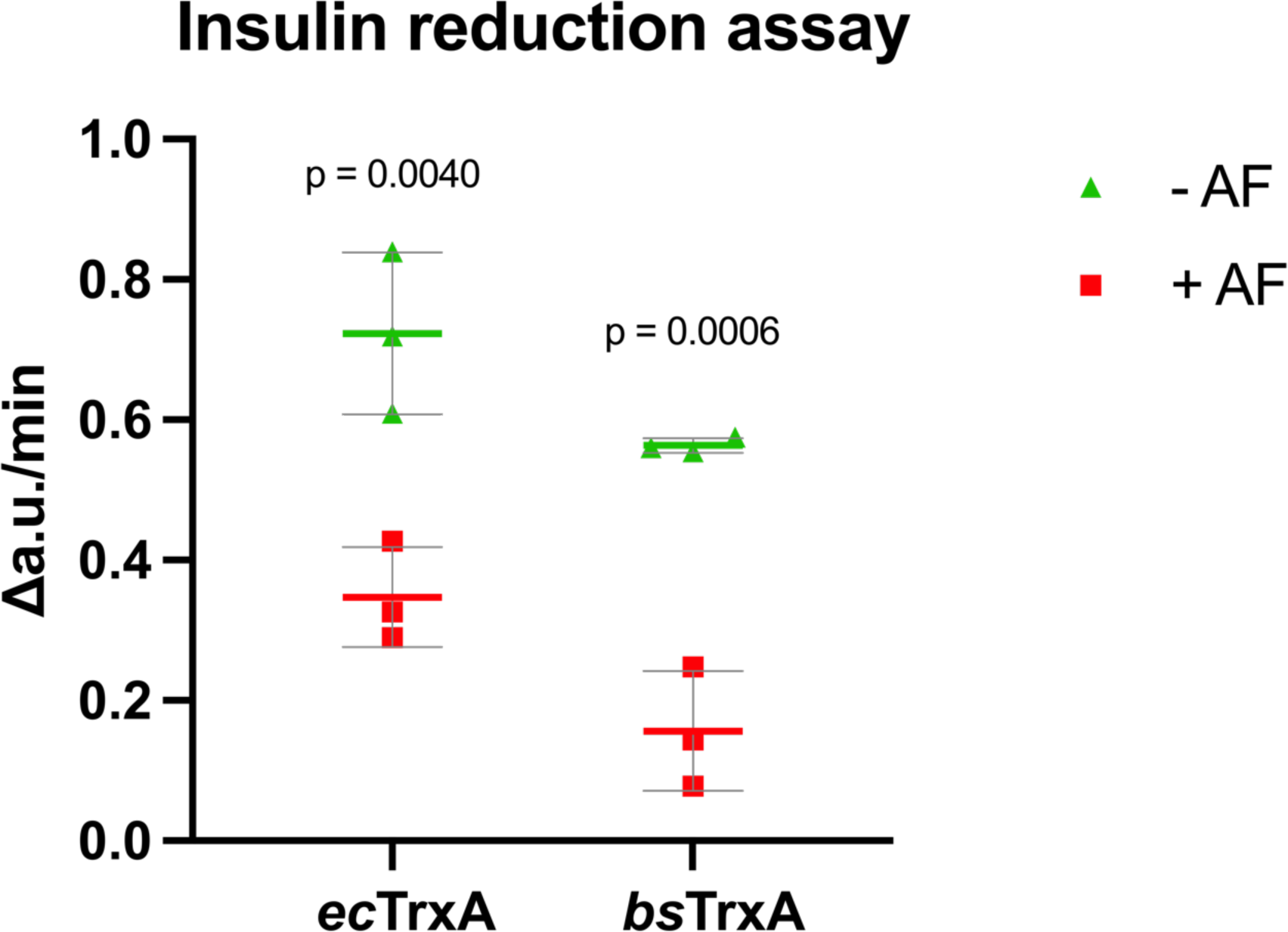
FITC-Insulin reduction by bacterial thioredoxins. Insulin reduction velocity of *ec*TrxA *(E. coli)* and *bs*TrxA (*B. subtilis*) is shown. 1 µM of purified TrxAs was added to 10 μg/ml FITC-labeled insulin. 1 µM of auranofin (AF) was added as inhibitor. Turnover was followed for 1 hour and the maximal velocity over 250 s was plotted. Mean, SD, and individual data points obtained from independent triplicates are shown. Significance (p-value) calculated using student’s T-Test. For raw data refer to supplementary material S2.

### Auranofin induces a modification that mimics a disulfide bond but is still reactive towards iodoacetamide (IAM)

Due to gold’s propensity to react with thiol sulfur, we hypothesized that the site of auranofin-binding in TrxA are the cysteines in the hallmark active site CXXC motif. In order to observe auranofin’s interaction with cysteines directly, we used roGFP2 as a molecular probe. roGFP2 is a genetically engineered variant of green fluorescent protein, which contains two cysteines that can be oxidized to form a disulfide bond (Aller et al., 2013). The fluorescence behavior of roGFP2 is contingent on the redox state of these two cysteines, and, based on the ratio of two particular bands in roGFP2’s excitation spectrum, its redox state can be determined. Thus, we suspected that roGFP2 could be a useful probe to monitor the interaction between auranofin and protein thiols. And indeed, adding auranofin to roGFP2 at a 1:10 molar excess led to a rapid spectral change (Fig. 3, a and b), highly similar to the spectral change observed when adding the well-characterized thiol-oxidant AT-2 (Fig. 3, c). Like the change induced by AT-2, it is reversible by the thiol reductant DTT (Fig. 3, b and c). We thus hypothesized that auranofin introduces a disulfide bond in roGFP2. To confirm this direct interaction, we pre-treated roGFP2 with iodoacetamide. Iodoacetamide is highly effective in blocking reduced thiols by carbamidomethylation. This “locks” roGFP2 in a pseudo-reduced state (Fig. 3 d), which no longer reacts with the thiol-oxidant AT-2 (Fig. 3 d). Similarly, auranofin can no longer change the fluorescence properties of iodoacetamide-treated roGFP2 (Fig. 3 d), confirming that, like the thiol oxidant AT-2, auranofin requires free thiols to react with roGFP2. While iodoacetamide reacts with free thiols, it cannot react with thiols engaged in a disulfide bond. Adding iodoacetamide to disulfide bond-containing roGFP2, thus, does not change its fluorescent properties (Fig. 3 e). Surprisingly, adding iodoacetamide to auranofin-treated roGFP2 significantly changes its fluorescent properties back to that of its pseudo-reduced carbamidomethylated form (Fig. 3 f). Taken together, this shows that the chemical alteration induced in roGFP2 by auranofin mimics a disulfide bond based on roGFP2’s spectral properties, but is chemically distinct, as it still possesses the ability to react with iodoacetamide.

**Figure 3.**
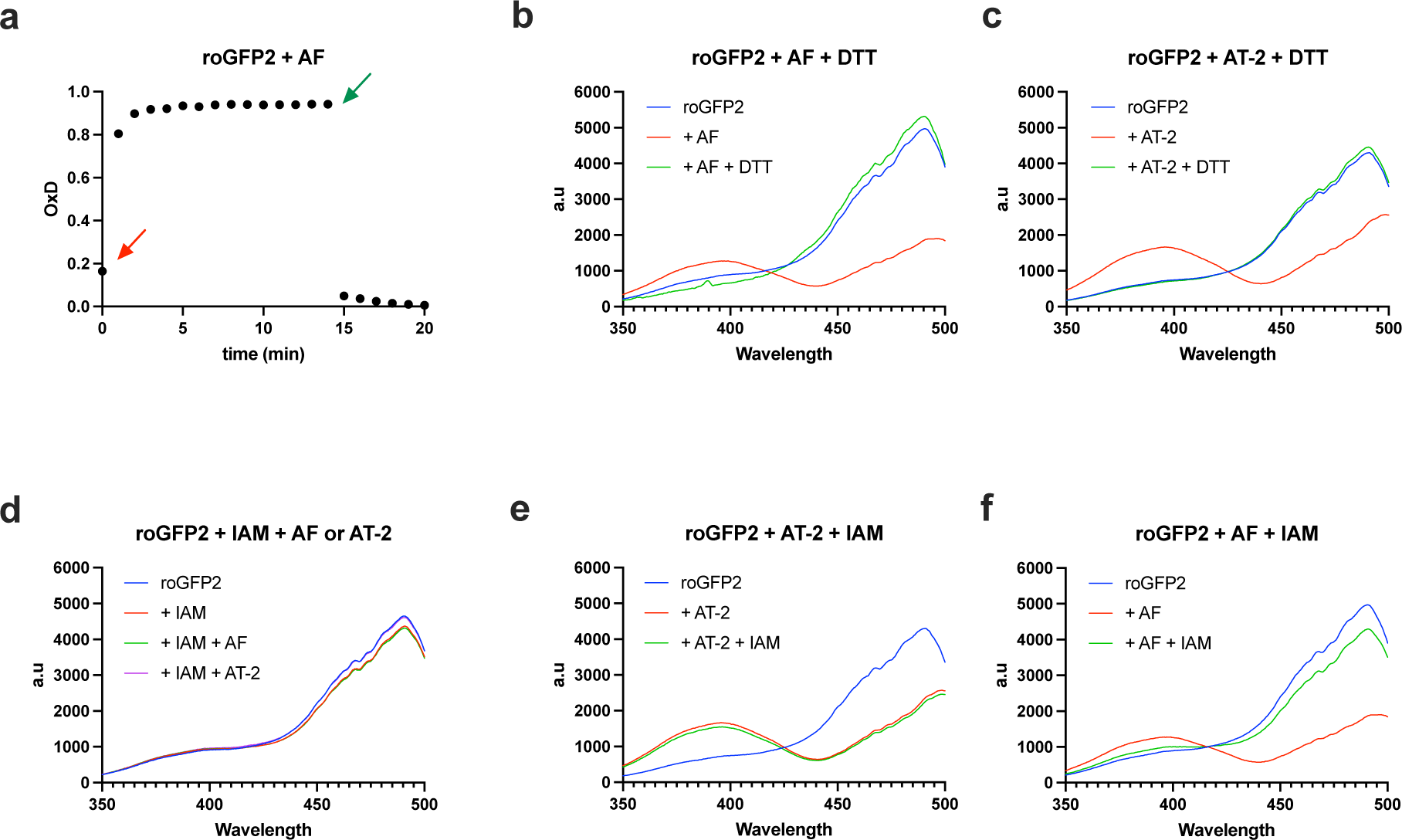
Changes in fluorescent spectra of roGFP2 treated with auranofin, AT-2, and the thiol-blocking agent iodoacetamide. **(a)** OxD of reduced roGFP2, treated with a 10-fold molar excess of auranofin (AF, indicated by a red arrow) and re-reduced with a 100-fold excess of DTT (indicated by a green arrow), demonstrating reversibility. **(b)** Spectral endpoints of the data presented in **(a)**. **(c)** A similar observation could be made when reduced roGFP2 was treated with a 10-fold molar excess of AT-2 and then re-reduced with a 100-fold molar excess of DTT. **(d)** Reduced roGFP2 was treated with a 1000-fold molar excess of iodoacetamide (IAM) to block all free thiols, and both AT-2 and auranofin (AF) were no longer able to induce a spectral change. **(e)** roGFP2 treated with a 10-fold molar excess of AT-2 was no longer susceptible to the addition of a 1000-fold molar excess of iodoacetamide (IAM). **(f)** However, roGFP2 treated with a 10-fold molar excess of auranofin (AF) returned to its pseudo-reduced state when treated with a 1000-fold molar excess of iodoacetamide (IAM). **(b-f)** Representative data shown, experiments were performed in independent duplicates **(b, c)** or triplicates **(d, e, f)**. Spectra of untreated roGFP2 in **(b)** and **(f)** are identical, as they were derived from the same experiment day and are repeated for illustrative purposes. The same applies to roGFP2 and AT-2-treated roGFP2 in panels **(c)** and **(e)**. For underlying data please refer to supplementary material S3.

### Auranofin forms a gold adduct in cysteine-containing proteins

The above data using roGFP2 showed that auranofin induced a thiol-modification similar to, but clearly distinct from, a disulfide bond. We thus speculated that the gold atom in auranofin could form a direct adduct with thiol pairs present in proteins. We used inductively coupled plasma-optical emission spectrometry (ICP-OES) to determine if gold is bound directly to cysteine thiols in roGFP2. In our experiment, we treated roGFP2 with auranofin and removed excess auranofin by gel filtration. Fluorescence spectrometry confirmed the successful modification of roGFP2 (Fig. 4, a), and the auranofin-treated roGFP2 protein contained Au in a 1:0.7 molar ratio, consistent with an equimolar modification (Fig 4b). roGFP2 pre-treated with iodoacetamide before treatment with auranofin did not contain gold (Fig 4b), in line with the spectral properties recorded (Fig. 4a). Treatment of auranofin-modified roGFP2 with iodoacetamide led to a substantial loss of gold, again in line with our previous data and the spectra recorded (Fig 3f and 4a). We thus concluded that auranofin-treated roGFP2 forms a 1 to 1 complex with Au, most likely as an S-Au-S bridge between the two structurally adjacent cysteines in roGFP2.

**Figure 4.**
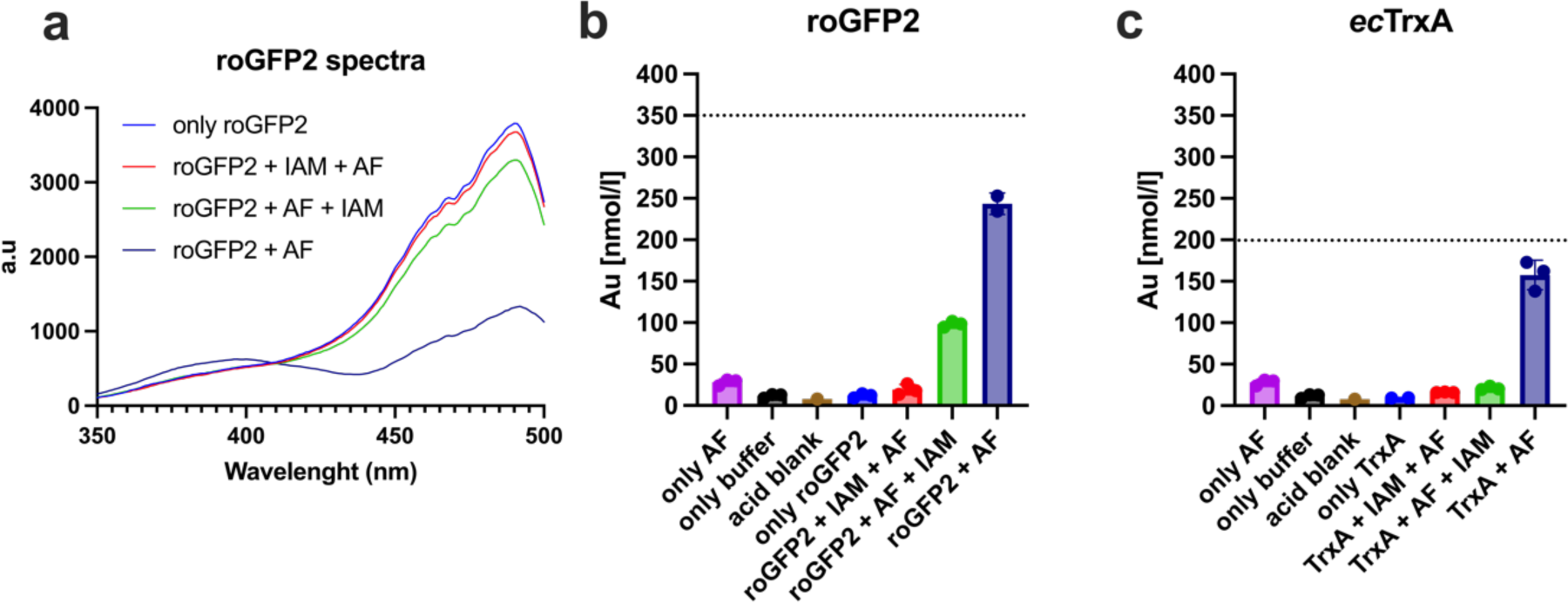
Au binding to cysteine-containing proteins. **(a)** Fluorescence spectra of roGFP2 samples subjected to ICP-OES analysis. ICP-OES analysis of **(b)** roGFP2 and **(c)** *ec*TrxA treated with auranofin (AF) and auranofin combined with iodoacetamide (IAM) in the order indicated (+IAM +AF: iodoacetamide first, then auranofin, +AF + IAM: auranofin first, then iodoacetamide). Proteins were subjected to a 10-fold molar excess of auranofin and a 1000-fold molar excess of iodoacetamide, where indicated. Excess auranofin was removed by gel-filtration. In the solutions analyzed, roGFP2 had a concentration of 350 nM **(b)**, while *ec*TrxA was present at a concentration of 200 nM **(c)** (indicated by dashed lines). Auranofin, in the absence of protein, was subjected to gel-filtration to control for carry over of gold (only AF), buffer without protein or auranofin (only buffer) and the analytical grade nitric acid used (acid blank) were analyzed as contamination controls. Controls are displayed in both panels **(b, c)** for illustrative purposes. Underlying fluorescence and ICP-OES data can be found in supplementary material S4.

Next, we asked ourselves if this was a peculiar property of roGFP2, or if this modification can also be found in TrxA, as we made our initial observations with this protein. Thus, we performed a similar experiment using *ec*TrxA. The ICP-OES data obtained with *ec*TrxA parallels the data obtained with roGFP2: auranofin treatment led to a 1:0.8 stoichiometry between protein and Au, which could be prevented by prior blocking of *ec*TrxA’s cysteines with iodoacetamide. More importantly, Au binding could be reversed by a subsequent reaction with iodoacetamide, confirming that *ec*TrxA, too, does not form a disulfide bond but a disulfide bond-mimicking Au adduct, which explains its inhibition by auranofin (Fig. 4 c).

### Auranofin-induced S-Au-S bonds in proteins are a substrate for thioredoxin

Next, we were wondering if the disulfide-mimicking thiol modification in proteins is, in principle, reversible by the antioxidant machinery available to a living cell. Thus, we treated auranofin-modified roGFP2 with rising concentrations of the thiol-reductase thioredoxin (Fig. 5). When reduced TrxA from *E. coli* was added to auranofin-modified roGFP2, we were able to monitor roGFP2’s fluorescence properties return to those of reduced roGFP2, suggesting that the gold-adduct in auranofin-treated proteins acts as a direct substrate of the thiol-disulfide reductase thioredoxin. Presumably, like a disulfide bond in a thioredoxin substrate protein, the S-Au-S bridge is transferred from the substrate protein onto thioredoxin.

**Figure 5.**
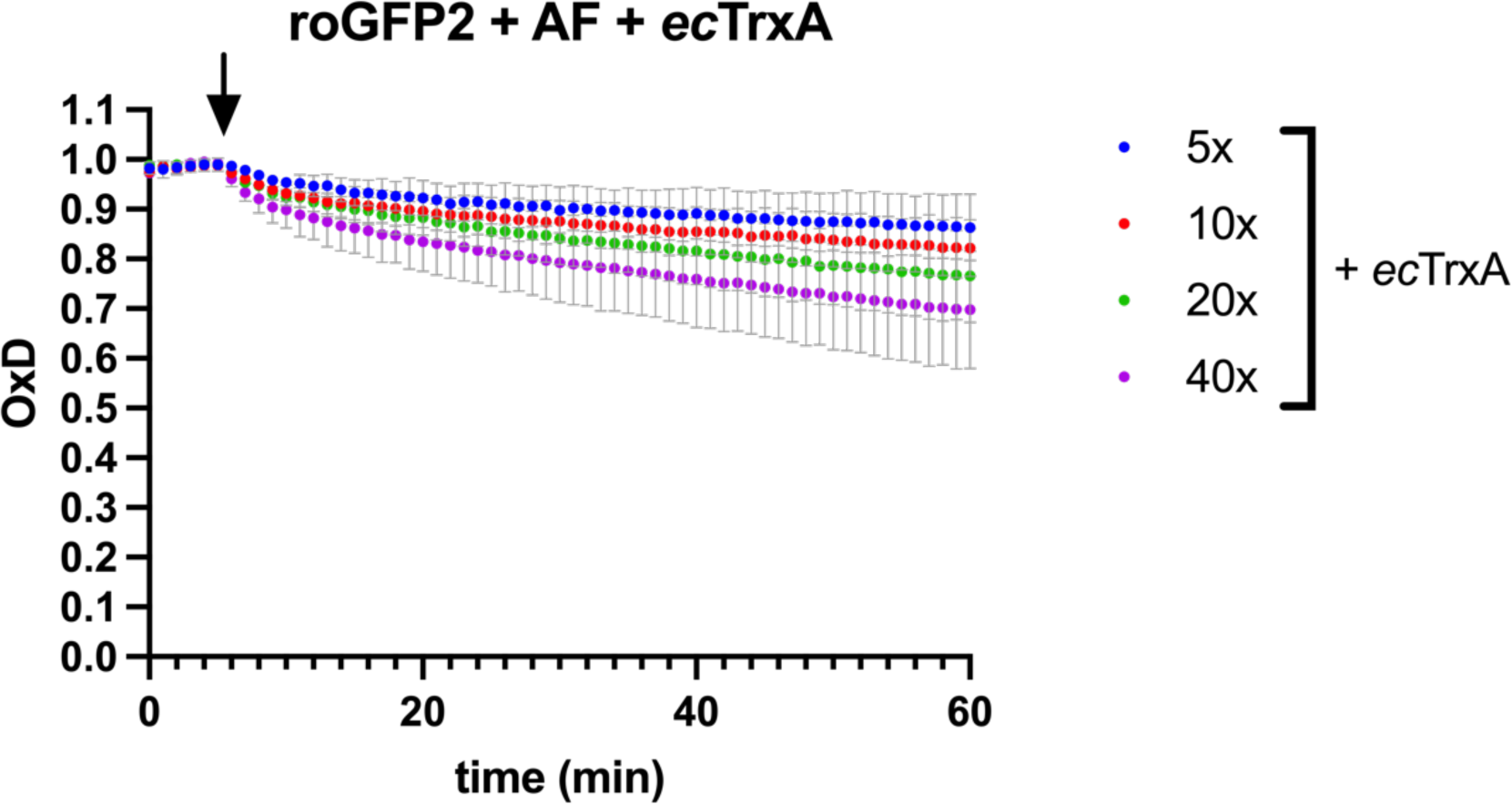
removal of S-Au-S adducts from roGFP2 recovery with *ec*TrxA. The oxidation degree (OxD) of roGFP2 treated with auranofin was evaluated fluorometrically. After stabilization of the signal, *ec*TrxA was added to the assay at the molar excess indicated, and OxD was monitored for a total of 1h. An arrow indicates the time point when *ec*TrxA was added. Experiments were performed in duplicates, average and range are displayed. The underlying sprectral data can be found in supplementary material S5.

## DISCUSSION

Here, we present a series of experiments elucidating the interaction between auranofin and cysteine pair-containing proteins, namely bacterial thioredoxin from *E. coli* and *B. subtilis*, and the thiol-redox probe protein roGFP2. Our data suggests that, in addition to inhibiting bacterial thioredoxin reductase TrxB, auranofin readily reacts with cysteine pairs present in thioredoxin reductase’s substrate thioredoxin and other proteins. Auranofin should, therefore, not be considered a specific inhibitor of thioredoxin reductase in bacteria.

In reconstituted bacterial thioredoxin systems, we observed K_m_ values of TrxB for TrxA in *E. coli* of 1.35 ± 0.23 µM, similar to values found by others ranging from 2.8 to 3.68 µM (Lennon & Williams, 1997; Prongay et al., 1989; Williams, 1976). In *B. subtilis,* the K_m_ was 1.72 ± 0.16 µM, comparable to a value reported elsewhere with a K_m_ of 4.22 ± 0.58 µM (Zheng et al., 2019). With these Michaelis-Menten kinetics established, we added varying concentrations of auranofin to our assays. And while auranofin’s inhibitory effect was concentration-dependent, our data did not conform to a typical inhibition process, preventing us from determining auranofin’s K_i_ and from obtaining meaningful insights into the mechanism of inhibition. Instead, thioredoxin itself seemed to attenuate TrxB inhibition in a concentration-dependent manner, with significant inhibition of TrxB observed when auranofin concentrations exceeded the concentration of TrxA.

In contrast, Owings *et al*. reported an IC_50_ of 88 nM for auranofin for *Helicobacter pylorii* TrxB in the presence of *E. coli* TrxA (Owings et al., 2016) and Harbut *et al*. reported IC_50_ values of 63 nM and 90 nM for thioredoxin reductases TrxB2 from *Mycobacterium tuberculosis* and TrxB from *Staphylococcus aureus* repectively, both in the presence of *M. tuberculosis* thioredoxin TrxC (Harbut et al., 2015). Owings *et al*. used TrxA at a concentration of 1 µM, well above their reported IC_50_ of 88 nM for auranofin, and Harbut *et al*. used TrxC at even higher concentrations of up to 50 µM, findings that seemingly contradict our observations: in our experiments, no or very little inhibition was observed when auranofin was used at concentrations below the concentration of thioredoxin. However, both groups did not preincubate thioredoxin reductase, NADPH, and auranofin with thioredoxin. Rather, they initiated the reaction by the addition of thioredoxin or thioredoxin, NADPH and DTNB. This is a sensible approach, when working under the assumption that auranofin exclusively interacts with TrxB. But should there be an additional formation of a thioredoxin-Au adduct, the transfer reaction of the Au adduct in TrxB to thioredoxin might not have been in equilibrium at the beginning of their enzymatic assay, leading to inhibition at the outset of the reaction which is then diminished over time.

Our speculation that auranofin interacts with TrxA was confirmed in thioredoxin activity assays determining TrxA’s ability to reduce FITC-labeled insulin. Both *ec*TrxA’s and *bs*TrxA’s ability to reduce insulin was significantly inhibited in the presence of auranofin. It is, therefore, tempting to speculate that TrxB inhibition is not the most relevant aspect of auranofin’s antibacterial mode of action, given that proteomic studies suggest that in *E. coli* and *B. subtilis,* the molar concentration of TrxA is 4.6 and 17.8 times higher than the concentration of TrxB, respectively (Muntel et al., 2014; Wiśniewski & Rakus, 2014). Our and others’ results of the comparatively similar susceptibility of mutants lacking TrxB and other components of the thiol-disulfide oxidoreductase machinery towards auranofin when compared to the wild type support this reasoning (Harbut et al., 2015; Thangamani et al., 2016). Given the fact that cells of higher eukaryotes also contain high amounts of cysteine pair-containing proteins (proteomic studies suggest a thioredoxin concentration 26.0 times higher than the concentration of thioredoxin reductase 1 in a commonly used human cell line (Beck et al., 2011)), thioredoxin reductase might not be the exclusive target in higher eukaryotes, either.

Using the engineered redox probe protein roGFP2 allowed us to directly observe the interaction of auranofin with cysteine thiols in a protein. Exposing reduced roGFP2 to auranofin *in vitro* led to a spectral change similar to the change induced by the well-established thiol oxidant AT-2. Like the disulfide bond generated by AT-2, the auranofin-induced modification was stable over time and reversible by thiol reductants such as DTT, and prior blocking of thiol residues with iodoacetamide prevents the alteration of roGFP2’s excitation spectrum. However, unlike a disulfide bond, the change in the spectrum can be reversed by the addition of iodoacetamide.

This indicated to us a direct interaction between gold(I) and the cysteines in roGFP2. ICP-OES revealed that gold is bound in an approximately 1:1 molar ratio to auranofin-treated roGFP2, and prior treatment with iodoacetamide prevents this gold binding. Similarly, iodoacetamide treatment after auranofin treatment significantly diminishes the amount of gold bound to roGFP2. All our data point to a direct interaction between the cysteine pair in roGFP2 and the gold ion in auranofin. Performing the same experiment with TrxA revealed the same pattern: an approximately 1:1 molar ratio of gold to protein, inhibition of gold binding by prior blocking of TrxA’s cysteines with iodoacetamide, and release of gold by iodoacetamide treatment of gold-containing TrxA. We thus concluded that these proteins form S-Au-S bonds, which functionally mimic disulfide bonds, as evidenced by the spectral change in roGFP2 and the inhibtion of TrxA’s reductase activity.

The observed reactivity of these protein S-Au-S bonds with iodoacetamide has implications on the interpretation of data obtained with methodologies typically used to determine the modification state of protein cysteines: as an example, recent studies have looked at the effect of auranofin on protein cysteines *in vivo* in redox proteomic studies (Saei et al. 2020; Chiappetta et al. 2022; Abu Hariri et al. 2022). These studies employed alkylating agents, such as iodoacetamide, iodoacetamide-based probes or N-ethylmaleimide to block “unmodified” cysteines. However, considering our findings, S-Au-S adducts that could have formed, would have probably reacted during the blocking step with these alkylating agents and thus would appear as unmodified cysteines in proteomic screenings. If one assumes, that S-Au-S bond formation in a protein has functional consequences (as evidenced by the inactivation of thioredoxin that we observed), redox proteomic studies using alkylating agents to determine cysteine modifications might underestimate the effect of auranofin on the functionality of cysteine-containing proteins.

The spectral change of roGFP2 exposed to auranofin observed here in vitro was also observed in vivo in bacterial cells expressing roGFP2 exposed to auranofin (Barse et al., 2023). The rate of spectral change in Gram-positive bacteria, which lack the low-molecular-weight thiol glutathione, was similar to the rate observed here in vitro, indicating that S-Au-S adducts also form in living cells, at a rate similar to the rate found in our *in vitro* findings.

Gold bound to a cysteine pair can be relocated from one reduced cysteine pair to another within the thiol-disulfide oxidoreductase thioredoxin glutathione reductase TGR (Angelucci et al., 2009; Saccoccia et al., 2014). We suspect that such a transfer, but between proteins, is the reason why inhibition of TrxB by auranofin is abrogated in the presence of high concentrations of TrxA, a mechanism suggested by Saccoccia et al. (Saccoccia et al., 2014). In line, we show that thioredoxin is indeed able to remove the spectrum-changing gold-adduct from roGFP2. These data suggest a potential protective role of the thioredoxin system against auranofin-induced stress. However, it remains unclear if the thioredoxin S-Au-S gold adduct is a dead end or if other systems in the cell can restore thioredoxin to its free thiol form. Short of reduction of Au(I) to its elemental form, only transfer to another thiol couple comes to mind as a thioredoxin-restoring mechanism. Here, low molecular weight thiols, such as glutathione, could play a major role. Indeed, we found that the presence of high concentrations of glutathione is one of the factors contributing most strongly to the low susceptibility of the Gram-negative bacterium *E. coli* to auranofin when compared to the Gram-positive bacterium *B. subtilis* (Barse et al., 2023).

Overall, our data suggests that thioredoxin reductase (TrxB) is not the exclusive target of auranofin in bacteria. Instead, thiol pairs are directly attacked at rates that imply S-Au-S bond formation as an integral part of auranofin’s antibacterial mode of action. Such a revised model of auranofin’s mode of action is consistent with a large extent of the data stated in the literature concerning auranofin’s effects *in vitro* and *in vivo*, also in higher eukaryotes. More importantly, it resolves the paradoxical nature of some of these observations, namely the many reports of IC_50_ or K_i_ values for an inhibitor that is presumably irreversible in nature: the transfer of gold(I) adducts from thioredoxin reductase to its substrates could be misinterpreted as reversible binding of the inhibitor. As long as the auranofin concentration in an assay is too low for all thiol pairs present to form S-Au-S bonds, residual thioredoxin reductase activity could potentially be observed, despite the irreversible nature of the actual inhibition reaction.

## AUTHOR CONTRIBUTIONS

L.Q.B and L.I.L. conceived and coordinated the study. N.L. purified all proteins used in this study. L.Q.B. performed experiments and P.D. ran the ICP-OES experiment and curated the associated data. J.B. and U.K. consulted on and contributed to analyzing and interpreting the results. L.Q.B and L.I.L. analyzed the data and wrote the manuscript, J.B., U.K., and P.D. edited the manuscript. All authors approved the final version of the manuscript.

## ACKNOWLEDGMENTS

L.I.L., J.E.B., and U.K. acknowledge funding from the German Research Foundation (DFG) through Research Training Group 2341 “Microbial Substrate Conversion (MiCon)”. L.I.L. received additional funding through the InnovationsFoRUM Host-Microbe-Interaction IF-018-22-TP8.

We thank Melissa Vázquez-Hernándes (Applied Microbiology, Faculty of Biology and Biotechnology, Ruhr University Bochum) for helping with the ICP-OES sample preparation. We thank Ingo Ott, Petra Lippmann and Pia Schneeberg (Institute of Medicinal and Pharmaceutical Chemistry, Technische Universität Braunschweig) for consulting on the ICP-OES sample preparation and analysis.

